# RlmQ: A Newly Discovered rRNA Modification Enzyme Bridging RNA Modification and Virulence Traits in *Staphylococcus aureus*

**DOI:** 10.1101/2023.09.27.559750

**Authors:** Roberto Bahena-Ceron, Chloe Teixeira, Jose R Jaramillo Ponce, Philippe Wolff, Florence Couzon, Pauline François, Bruno Klaholz, François Vandenesch, Pascale Romby, Karen Moreau, Stefano Marzi

**Affiliations:** Université de Strasbourg, CNRS, Architecture et Réactivité de l’ARN, UPR9002, F-67000 Strasbourg, France; CIRI, Centre International de Recherche en Infectiologie, Université de Lyon, Inserm U1111, Université Claude Bernard Lyon 1, CNRS UMR5308, ENS de Lyon, Lyon, France; Centre for Integrative Biology, Department of Integrated Structural Biology, IGBMC, 1 rue Laurent Fries, Illkirch, France; CNRS UMR 7104, Illkirch, France; Inserm U964, Illkirch, France; Université de Strasbourg, Strasbourg, France; Institut des agents infectieux, Hospices Civils de Lyon, Lyon, France; Centre National de Référence des Staphylocoques, Hospices Civils de Lyon, Lyon, France

**Keywords:** m^7^G in 23S rRNA, RNA modification, methyltransferase, tRNA accommodation, *Staphylococcus aureus* virulence

## Abstract

rRNA modifications play crucial roles in fine-tuning the delicate balance between translation speed and accuracy, yet the underlying mechanisms remain elusive. Comparative analysis of the ribosomal RNA modifications in taxonomically distant bacteria could help define their general as well as species-specific roles. In this study, we identified a new methyltransferase, RlmQ, in *Staphylococcus aureus* responsible for the Gram-positive specific m^7^G2601, which is not modified in *E. coli* (G2574). We also demonstrate the absence of methylation on C1989, equivalent to *E. coli* C1962, which is methylated at position 5 by the Gram-negative specific RlmI methyltransferase, a paralogue of RlmQ. Both modifications (*S. aureus* m^7^G2601 and *E. coli* m^5^C1962) are situated within the same tRNA accommodation corridor, hinting at a potential shared function in translation. Inactivation of *S. aureus rlmQ* causes the loss of methylation at G2601 and significantly impacts growth, cytotoxicity, and biofilm formation. These findings unravel the intricate connections between rRNA modifications, translation, and virulence in pathogenic Gram-positive bacteria.

## INTRODUCTION

Ribosomal RNA (rRNA) is an essential component of the ribosome, the cellular machinery responsible for protein synthesis. It plays the most critical functional roles, assisting mRNA decoding, tRNA movements and peptide bond formation (Nissen et al. 2000; Ogle et al. 2001; Demeshkina et al. 2012; Noller et al. 2017). Over the years, extensive studies have identified various post-transcriptional modifications that adorn rRNA molecules. These modifications, which are added during ribosome assembly (Siibak and Remme 2010), participate in the balance between translation speed and accuracy. Indeed, *in vitro* reconstituted *Escherichia coli* ribosomes lacking rRNA modifications were severely defective in catalytic activity (Green and Noller 1996) and the ribosome assembly was also altered (Cunningham et al. 1991). Numerous studies showed that the loss of rRNA modifications perturbs the active site structures in bacterial (Desaulniers et al. 2008; Demirci et al. 2010) and eukaryotic/human ribosomes (Natchiar et al. 2017), and causes altered rates and accuracy of translation (Liang et al. 2007).

In bacterial ribosomes, three primary types of rRNA modifications exist: pseudouridine (Ψ), methylation of the 2’-hydroxyl group of riboses (Nm), and methylation of the base (mN) (Decatur and Fournier 2002). While the precise roles of numerous modifications remain undetermined, they impart distinct characteristics to the nucleotides. For example, they can changes the hydrogen bond capacities (Ψ) or decreased stacking of bases (Dihydrouridine, D), impart structural rigidity (Ψ and Nm) or flexibility (mN) to both single and double-stranded regions, potentially affecting hydrogen bonding (Arnez and Steitz 1994; Charette and Gray 2000). These modifications are concentrated in highly conserved and dynamic regions crucial for ribosomal functions including decoding, peptidyl transfer, the binding sites of A- and P-site tRNAs, the peptide exit tunnel, and the inter-subunit bridges (Golubev et al. 2020; Watson et al. 2020).

The conservation of the position and chemical nature of the modified nucleotide across bacterial taxa has been a subject of growing interest. Several modifications appear to be strictly conserved, while others vary among bacteria. For instance, within the large ribosomal subunit of *Escherichia coli* ribosome, Gm2251 and Um2552 are situated in the P- and A-loops, respectively, where they interact with the CCA end of tRNAs in the P- and A-sites. While Gm2251 is conserved across the three domains of life (Sergeeva et al. 2015; Natchiar et al. 2017), Um2552, added by RlmE methyltransferase, is present in many bacteria, except for certain Bacillus species like *Bacillus subtilis* and *Bacillus stearothermophilus*. Instead, in these bacterial species, a different methyltransferase, RlmP, methylates G2553 during 50S biogenesis (Hansen et al. 2002; Roovers et al. 2022). In *rlmE*-deficient *E. coli* mutant strain, the absence of Um2552 modification, which decreases the flexibility of adjacent nucleotides, leads to a delay in 50S subunit maturation, to a slower subunit association, and translocation rate (Wang et al. 2020). Translation with unmethylated U2552 in *E. coli* appears to be more accurate (Widerak et al. 2005) and to proceed at a reduced rate (Caldas et al. 2000). These results suggested that a certain level of recoding provided by the methylation can be accepted to favor higher translation speed important for cellular physiology. Numerous rRNA modifications have also been observed along the tRNA transition pathway, both on the 30S and the 50S subunits (Watson et al. 2020). During early ribosome assembly, the methyltransferase RlmI methylates at position 5 of C1962 (Purta et al. 2008), which is located on helix H70 at the edge of its coaxial stacking with H71. Together with H69, which contains the conserved Ψ1911 (RluD), m^3^Ψ1915 (RluD, RlmH), and Ψ1917 (RluD), these rRNA helices provide a sliding support for the movements of tRNAs into the ribosome (Girodat et al. 2023). The *rlmI*-depletion in a mutant *E. coli* strain led to a small growth defect in competition experiments (Purta et al. 2008) or when translation is enhanced following the overexpression of an induced gene (Pletnev et al. 2020).

In the present study, we have identified a paralog of *E. coli* RlmI in *Staphylococcus aureus*. Unexpectedly, we find that the enzyme is responsible *in vivo* for the methylation at position 7 of G2601 located in helix H90 of the 23S rRNA. Importantly, this position is unmodified in *E. coli* (G2574) hinting at a specific function in *S. aureus*. Because we did not detect any methylation at C1989 of *S. aureus* 23S rRNA (equivalent to *E. coli* C1962 methylated at position 5), we have renamed this new methylase, RlmQ. Phenotypic analysis of the *rlmQ*-disruption mutant strain revealed a slightly slower doubling time in nutrient-poor media, a reduced cytotoxicity, and an increased biofilm production. In the following, we will also discuss evolutionary considerations of RlmQ and RlmI, two highly similar enzymes with distinct specificities for 23S methylation in distant bacterial species, and their biological consequences.

## RESULTS AND DISCUSSION

### *S. aureus* 23S rRNA contains a 7-methylguanosine at position 2601

Given the important roles of RNA modifications on bacterial physiology, we wondered whether the Gram-positive *S. aureus* bacteria harbor different sets of RNA modifications compared to the evolutionary distant Gram-negative bacteria *E. coli*. Here, we have focused the study on the modifications present in *S. aureus* 23S rRNA. After purification of the *S. aureus* ribosome from the HG001 strain, the identification of 23S rRNA modifications was done using RNase T1 digest followed by LC/MSMS mass spectrometry analysis. Among the various sequenced oligoribonucleotides, a fragment with a mass of 1948.2 Da (m/z 973.62) corresponded to the UAC[mG]CG>p sequence. Subsequent MSMS fragmentation revealed the presence of a methylation at G2601 located in H90 (corresponding to *E. coli* G2574; Figure 1A and 1B). To visualize the presence of the methylation and its position on the guanosine, we conducted an in-depth analysis of the high-resolution structure of *S. aureus* ribosome (Halfon et al. 2019) (map: EMD-10077; PDB: 6s0z; Figure 1C). Within this map, we identified a density that extends into position 7 within the purine ring, in favor of the presence of a 7-methylguanosine (m^7^G). Finally, the presence of m^7^G was validated by Primer Extension (PE) analysis, adapting a specialized protocol originally designed for genome-wide mapping of m^7^G (Marchand et al. 2018). The experiments showed a strong band corresponding to a specific termination of reverse transcription due to m^7^G in both the virulence-attenuated HG001 and the virulent USA300 (LAC) strains (Figure 1D). Interestingly, this modified position has also been reported in *Bacillus subtilis* (Hansen et al. 2002) suggesting that m^7^G2601 in the helix H90 region is conserved among firmicutes. *E. coli* ribosome structures suggest that H90 serves as a corridor to facilitate the accommodation of the aminoacylated CCA-end of tRNA (aa-CCA) into the A-site of the Peptidyl Transferase Center (PTC) (Whitford et al. 2010) (Figure 1E). Although the resolution of the cryo-EM structure of *S. aureus* 70S ribosome is lower than *E. coli* 70S and no A-site tRNA is present (Golubev et al. 2020), the positioning and structures of H90 and neighbor rRNA helices is highly similar. The accommodation process involves key conformational changes in both tRNA and ribosome that significantly impacts the rate of cognate tRNA selection (Rodnina and Wintermeyer 2001). A kinetic study combined with dynamic simulation has revealed the relatively slow nature of this process due to spontaneous and reversible accommodation attempts, akin to a stochastic trial-and-error mechanism, encompassing various parallel pathways (Whitford et al. 2010). Notably, the corridor traversed by the aa-CCA of the tRNA is nearby m^7^G2601 in *S. aureus*. Hence, it is tempting to propose that this methylation might be one of the key players which facilitates the accommodation process. More experiments will be required to assess the impact of the m^7^G modification on translation dynamics, especially its selectivity and functional importance for different aminoacyl-tRNAs.

**Figure 1.**
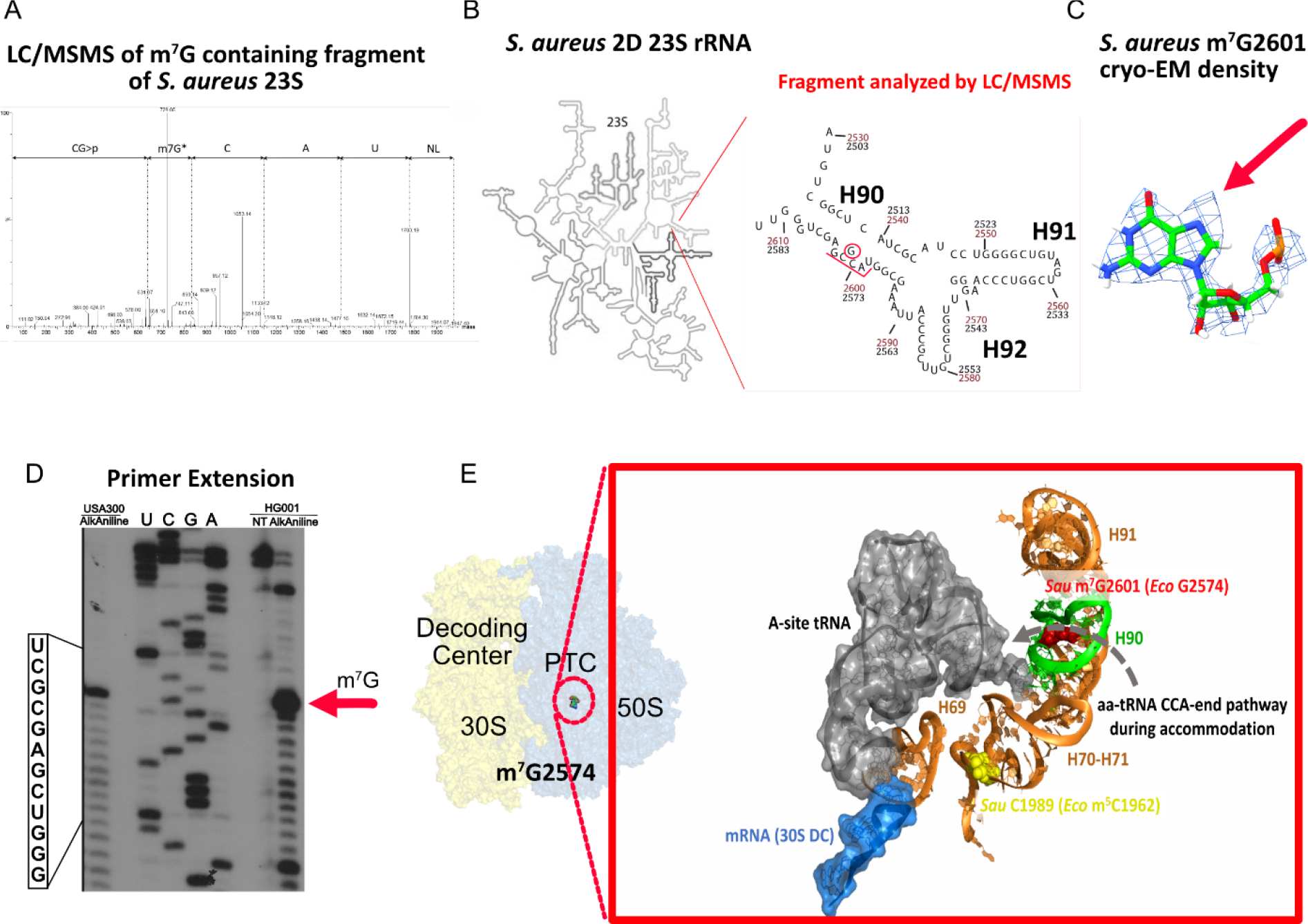
*S. aureus* ribosome contains m^7^G2610. A) Mass spectrometry analysis of the RNase T1 derived fragment of *S. aureus* HG001 23S rRNA region spanning from U2598 to G2603 (corresponding to *E. coli* U2571 to G2576). The fragment has a mass of 1948.2 Da (m/z 973.62) and carries the sequence UAC[mG]CG>p based on MSMS analysis. B) 2D representation of the entire *S. aureus* 23S rRNA, zooming on the region containing helix H90 with G2601 (corresponding to *E. coli* G2574). Red and black numbers indicate *S. aureus* and *E. coli* numbering, respectively. The red bar highlights the fragment analyzed by LC/MSMS. C) Structural analysis of G2601 from the deposited *S. aureus* 50S cryo-EM map at 2.3 Å (Halfon et al. 2019) (map: EMD-10077; PDB: 6s0z). The density around N7 indicates the presence of the methyl group, which is not modelled. D) Primer Extension analysis of HG001 and USA300 (LAC) *S. aureus* strains, including controls for the AlkAlanine treatment. AGCU lanes represent dideoxy sequencing, NT stands for the non-treated sample and the red arrow points to the reverse transcriptase stop due to the m^7^G modified nucleotide. E) Details of the accommodation corridor for aminoacyl-tRNA CCA-end on the *E. coli* ribosome structure, with the A-site tRNA (grey surface) in the accommodated state (Watson et al. 2020). The *E. coli* 23S rRNA helices (H69-H71, H90-H91) are shown: G2524 (red sphere), corresponding to m^7^G2601 in *S. aureus*, is located in H90 (green), while m^5^C1962 (yellow sphere), corresponding to C1989 in *S. aureus*, is in H70/H71 (bright orange). The mRNA (blue transparent surface) in the 30S decoding channel (DC) interacts with the anticodon of the A-site tRNA. *Sau*: *S. aureus*; *Eco*: *E. coli*.

### In quest of the enzyme responsible for m^**7**^**G2601**

To identify the enzyme responsible for m^7^G2601 in *S. aureus*, we have first performed a bioinformatic analysis, primarily focusing on sequence similarities through BlastP and sequence alignments. Our approach has involved three representative methyltransferases from *E. coli* and *B. subtilis*, which are known for their methylation activity on 16S rRNA (RsmG), 23S rRNA (RlmLK), and tRNA (TrmB). This investigation reveals seven potential m^7^G-methyltransferase genes in the *S. aureus* genome. To ascertain whether one of these enzymes was responsible for the presence of m^7^G2601, we systematically examined the PE profile of 23S rRNA prepared from the mutant strains carrying individual gene disruption for each candidate (Figure 2A). These strains were selected from the Nebraska Transposon Mutant Library collection and compared with the WT USA300 JE2 strain (Fey et al. 2013). The PE analysis showed that only the mutation at the locus SAUSA300_0981 in USA300 (HG001_00931 in HG001 strain) causes the loss of the specific RT pause at G2601. Complementation of the mutant strain transformed with a plasmid expressing the wild-type (WT) SAUSA300_0981 gene under the control of a constitutive mild promoter, restores the PE premature stop (Figure 2A). All in all, these experiments show that SAUSA300_0981 encodes the methyltransferase responsible for the 7-methylation of G2601. In accordance with the established bacterial nomenclature for rRNA methyltransferases, we have named this newly identified methyltransferase “RlmQ”.

**Figure 2.**
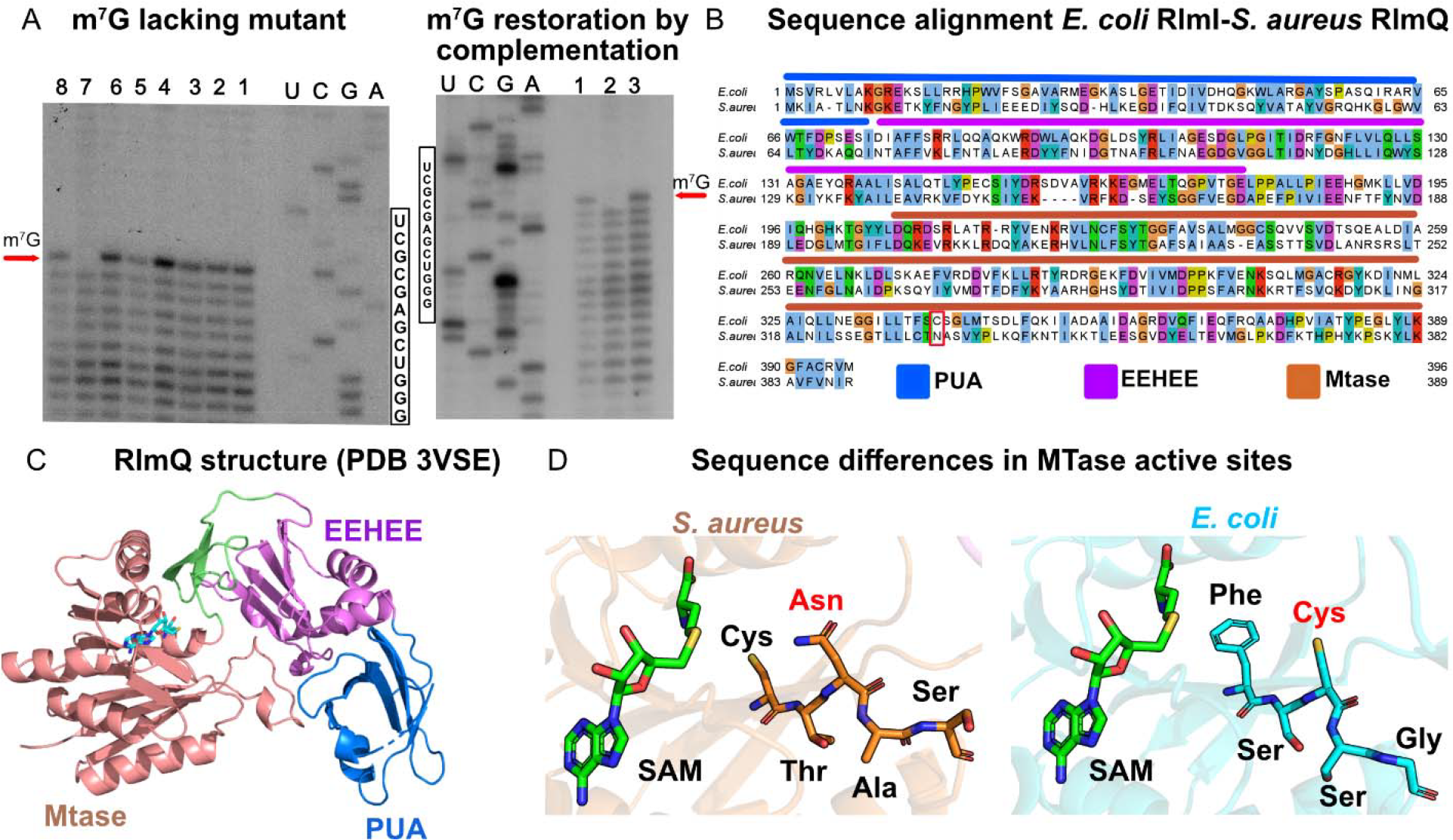
RlmQ is responsible for m^7^G2601. A) AlkAniline Primer Extension analysis of *S. aureus* 23S rRNA from USA300 JE2 WT and selected strains from the Nebraska collection. AGCU dideoxy sequencing lanes. Left: Lane 1, USA300 WT; lane 2, NE553; lane 3, NE1671; lane 4, NE1442; lane 5, NE189; lane 6, NE807; lane 7, NE1546 (corresponding to *rlm*Q-disruption mutant); lane 8, NE1337. Right: Analysis of m^7^G restoration. Lane 1, *S. aureus* USA300 Lac WT-pCN38:wP:TT; lane 2, NE1546-pCN38:wP:TT; lane 3, NEB1546-pCN38:wP:*rlm*Q:TT. Red arrows indicate m^7^G modification. B) Sequence alignment of *E. coli* RlmI and *S. aureus* RlmQ. PUA domain (blue bar over the sequence), EEHEE domain (magenta bar) and MTase domain (red bar) are shown by colored bars. The red box highlights the substitution of the conserved cysteine, typically found in m^5^C methyltransferases, by an asparagine in RlmQ. C) Crystal structure of *S. aureus* RlmQ, pdb files 3VSE (Kita et al. 2013). In blue the PUA domain, in magenta the EEHEE domain and in red the MTase domain. In cyan, the S-Adenosyl-Methionine cofactor. D) Comparison of the MTase active sites of *S. aureus* RlmQ (orange background structure) and *E. coli* RlmI (cyan background structure). While *E. coli* RlmI contains a cysteine (Cys) in the motif FS**C**SG, *S. aureus* RlmQ has an asparagine (Asn) in the motif CT**N**AS.

### *S. aureus* RlmQ and *E. coli* RlmI methyltransferases are paralogues with distinct activities

Based on its amino acid sequence, HG001_00931 (or SAUSA300_0981 in USA300) gene locus was previously annotated as RlmI (Caldelari et al. 2017), an RNA methyltransferase responsible for m^5^C1962 in 23S rRNA from *E. coli* (corresponding to *S. aureus* C1989) and various bacteria (Purta et al. 2008). Indeed, they share 46% sequence similarity and signatures for similar domain organization (Figure 2B). To perform a more detailed analysis of the similarities and differences between these two 23S rRNA methyltransferases, we conducted an extensive search in the PDB database using RlmQ as the reference (Holm et al. 2023). Remarkably, we identified the crystal structure of a *S. aureus* methyltransferase named SAV1081 (Kita et al. 2013), with no attributed function but which displayed 100% coverage and sequence similarity to RlmQ. This structure displays three main domains: a Rossman-Fold methyltransferase domain (MTase), the methyltransferase conserved RNA recognition module EEHEE domain, and the RNA binding domain PUA, which is responsible for the recognition of specific RNA sequences (Sunita et al. 2008) (Figure 2C). The EEHEE and MTase domains are interconnected by a conserved ß-hairpin structure, a feature shared with several other methyltransferases. Superposition of *E. coli* RlmI and *S. aureus* RlmQ structures (pdb files 3VSE for RlmQ and 3C0K for RlmI) shows a root mean square deviation (RMSD) of 1.37 Å, indicating a strong structural similarity. Both enzymes belong to the COG1092 family, with a well conserved S-Adenosyl-Methionine (AdoMet) binding site in the MTase domain. Additionally, their EEHEE domains exhibit a high degree of conservation, whereas their N-terminal PUA domains display the most pronounced divergence. Within the MTase active site, the principal distinction arises from a specific amino acid substitution: a catalytic cysteine (C) in RlmI is substituted by an asparagine (N) in RlmQ (Figure 2D). The presence of this cysteine in Gram-negative bacteria confers m^5^C RNA methyl transferase activity (Liu and Santi 2000; King and Redman 2002), while the presence of asparagine has been found in a separate subgroup of the COG1092 family in methylases mainly including Gram-positive members (Sunita et al. 2008).

To ascertain whether RlmQ was also involved in the modification of m^5^C1962, we conducted a meticulous analysis using mass spectrometry on the relevant 23S rRNA region (Figure 3A). The 23S rRNA, isolated from HG001 strain, was first hybridized to two specific oligonucleotides and digest with RNase H. This treatment led to a specific RNA fragment of 25 nucleotides, which includes rRNA helices H70 and H71 and C1962 (Figure 3B). After gel electrophoresis purification, the gel band containing the 25 nt long RNA fragment was further treated with RNase T1. Subsequent LC/MSMS analysis revealed the presence of a short fragment with a mass of 2249.3 Da corresponding to the unmodified sequence ACCCGp (Figure 3A). Comparison of the density maps from the reported high-resolution structures of *E. coli* and *S. aureus* ribosomes reveals clearly the presence of a methyl group at position 5 of C1962 in *E. coli* 23S while this methyl is clearly absent at position 5 of the equivalent C1989 in *S. aureus* (Figure 3C).

**Figure 3.**
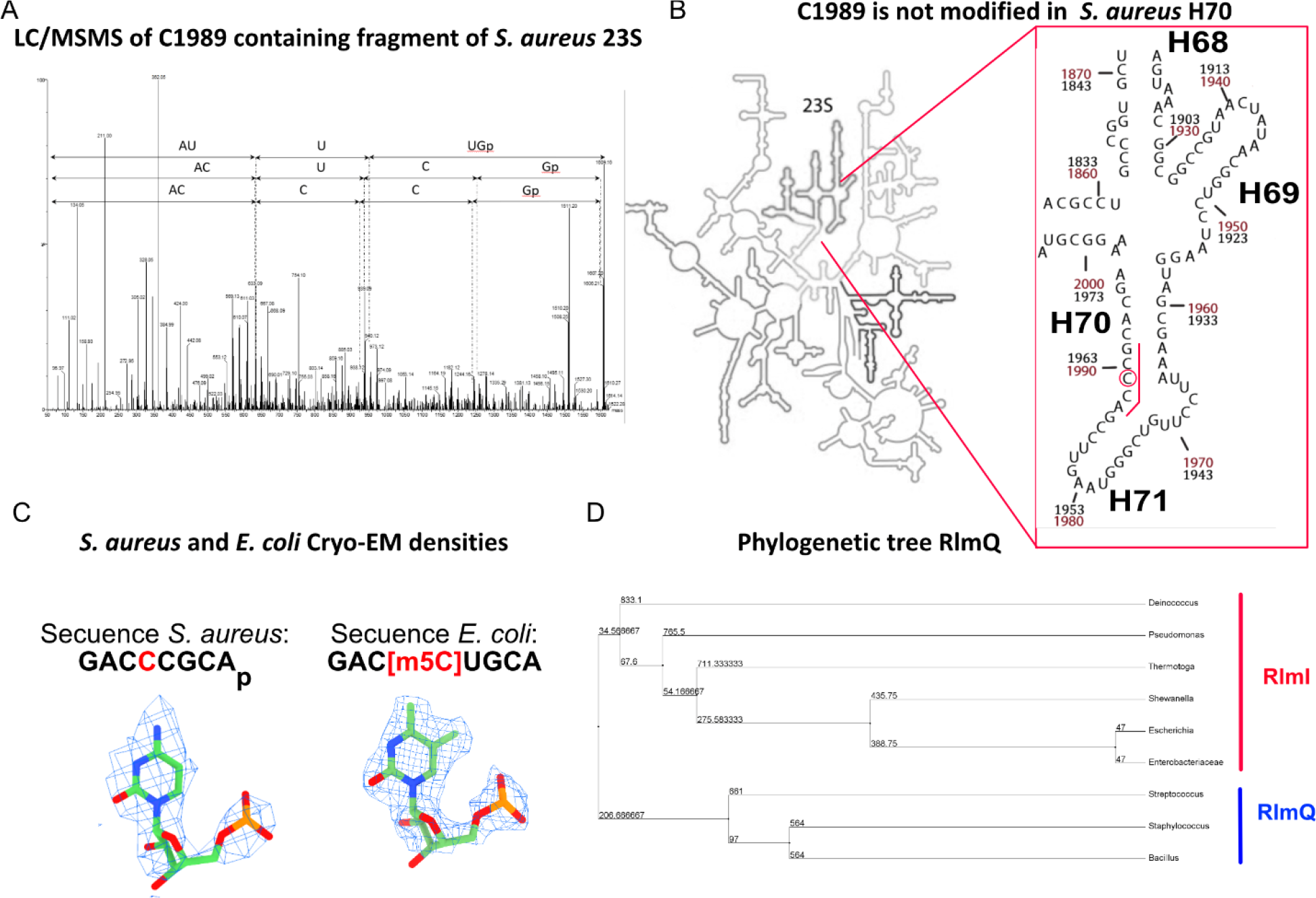
RlmQ evolution changes 23S methylation map. A) Detection of non-modified C1962 using mass spectrometry. The RNase H generated fragment including H70 and H71 of *S. aureus* 23S (from nucleotide A1975 to U2068) has been further digested with RNase T1 before analysis. MSMS spectra include the sequence ACCCGp showing methylation. B) 2D representation of the entire *S. aureus* 23S rRNA, zooming on the region containing H70 and H71, with C1962 (*E. coli* numbering, corresponding to *S. aureus* C1989). Red and black numbers indicate *S. aureus* and *E. coli* numbering, respectively. The red bar highlights the fragment analyzed by LC/MSMS. C) Structural analysis of C1962 from the deposited *S. aureus* 50S cryo-EM map at 2.3 Å (Halfon et al. 2019) (map: EMD-10077; PDB: 6s0z) and *E. coli* deposited 70S cryo-EM map at 2 Å (Watson et al. 2020) (map: EMD-22586; PDB: 7k00). While a clear density for methylation of position 5 on C1962 is observed in *E. coli* 23S allowing building this modification in the fitted model, no extra density is present on *S. aureus* 23S. D) Phylogenetic tree of RlmI-related family proteins. Sequences are denoted by their National Center for Biotechnology Information (NCBI) numbers. Node values represent statistical support for branches determined through bootstrap testing.

Taken together, we show *in vivo* that *S. aureus* RlmQ methylates the N7 position of G2601 (equivalent to G2574 in *E. coli*) while C1989 is not modified (equivalent to C1962 in *E. coli*). Because this methylation has also been observed in *B. subtilis*, we propose that Gram-positive bacteria share the same features. Conversely, in Gram-negative bacteria, RlmI is the enzyme which methylates C1962 at its position 5, while no existing modification was attributed to G2574 (Figure 3D). It is conceivable that these two paralogous enzymes may have originated from a gene duplication event and divergent evolution in these distant species. Interestingly, *E. coli* m^5^C1962 and *S. aureus* m^7^G2601 are both situated along the pathway taken by tRNAs during their transit into the ribosome (Watson et al. 2020) (Figure 1E). In this scenario, the tRNAs do not make direct contacts with helices H70 and H71, although these helices are in close proximity to the tRNA acceptor arms and variable loops. It is plausible that Gram-negative and Gram-positive ribosomes have evolved slightly different solutions to facilitate the smooth transition of tRNAs within the ribosome, both involving rRNA methylation, albeit targeting distinct residues.

### Phenotypic characterization of *S. aureus rlmQ* mutant

*S. aureus* is an opportunistic pathogen, which has evolved many strategies to regulate the synthesis of numerous virulence factors in response to the host, stress, and various environmental changes. To assess the impact of RlmQ enzyme modification on the physiology of *S. aureus*, the WT USA300 JE2 strain was compared to a mutant deficient for the *rlm*Q enzyme (*rlm*Q^-^) in various phenotypic tests including growth rate, biofilm, and cytotoxicity on human cells (Figure 4). To ensure that the differences observed were solely attributable to RlmQ, we have sequenced and analysed the genomes of the two strains. The presence of the transposon at position 448 of the *rlmQ* gene sequence (1173pb long) was validated and no other sequence changes in the genome was detected. The *rlm*Q^-^ mutant can therefore be considered as an isogenic mutant of JE2 strain where G2601 is not methylated (Figure 2A).

**Figure 4.**
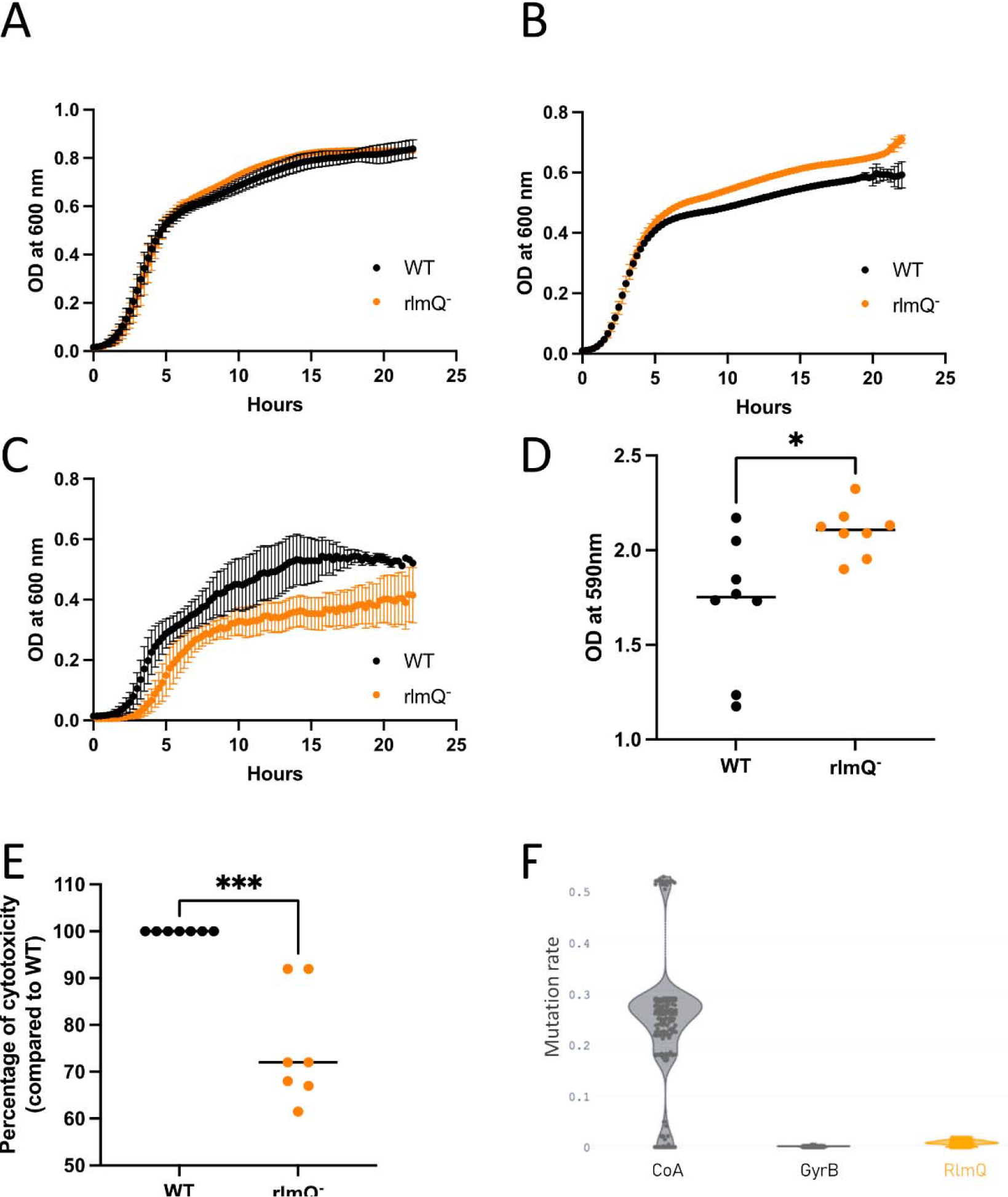
Phenotypic characterization of *S. aureus rlmQ* mutant. (A, B, C) Growth of the wild-type strain (WT) and the mutant inactivated for *rlm*Q (rlmQ^-^) was compared under different media and temperature conditions. Bacterial growth was measured during 24 h in BHI medium at 37°C (A), 42°C (B) and in RPMI medium at 37°C (C), n=2-6. (D) Biofilm was quantified by crystal violet staining extracellular matrix after 24 h of culture in BHI medium, n=8. (E) Bacterial surpernatant cytotoxicity was quantified by measuring propidium iodide incorporation into C5aR1-transfected U937 monocytes, n=7. Mann-Whitney tests performed: * = p-value < 0.05, ** = p-value < 0.01, *** = p-value < 0.001. F) Mutation rate of protein sequences from 301 clinical strains was estimated with a ratio: number of missense mutations divided by sequence match length.

We analyzed whether the mutation had an effect on *S. aureus* growth. Under optimal growth conditions (BHI, at 30°C and 37°C), no difference was observed between the WT and the *rlm*Q^-^ mutant strains (Figure 4A, supp Figure 1). This is in agreement with the fact that the polysome profiling in the WT and mutant strains remained identical in the two strains under these conditions of growth (supp Figure 2). These experiments also show that the maturation process of the ribosome is not significantly affected by the mutation. Only at 42°C in BHI, the *rlm*Q^-^ mutant reached a slightly higher OD at the stationary phase than the WT strain (Figure 4B). The two strains were cultured in a less rich environment (RPMI medium) closer to the conditions encountered during infection. Under these conditions, we observed a significant reduction in the growth of the mutant strain compared to the WT strain with a reduced doubling time at the exponential phase (*rlm*Q^-^ 65 min/WT 45 min), and a diminution of the maximal OD (*rlm*Q^-^ 0,38/WT 0,56) (Figure 4C). We then monitored the ability of *S. aureus* WT and mutant strains to form biofilm which play major roles in an infectious context. The amount of biofilm formed on plates by the *rlm*Q^-^ mutant was significantly greater than that of the WT strain (Figure 4D). Another important feature of bacterial-host interaction in infectious conditions is the cytotoxic activity of *S. aureus* against immune cells. The cytotoxic activity of the WT and mutant strains was assessed on human monocytes expressing the C5aR1 receptor targeted by two pore forming toxins, the Panton Valentine leucocidin (PVL) and the gamma toxin CB (HlgCB). The *rlm*Q^-^ mutant showed a significant reduction in cytotoxic activity compared with the WT strain (Figure 4E).

We finally addressed whether RlmQ is conserved among clinical infection isolates. To this end, the *rlm*Q gene was searched in the genomes of a collection of 301 well-characterized clinical *S. aureus* strains from previously published cohorts. The data showed that the *rlm*Q gene was detected in all 301 strains and had the same size in 298 strains (Figure 4F). Furthermore, the protein sequence was extremely well conserved with a mutation rate of 0,008 comparable to the highly conserved GyrB (MR=0,002) but much lower than that of coagulase (MR = 0263), which is known to be highly variable among strains. Using a Random Forest model based on horizontal oligonucleotide frequency, a conserved promoter sequence was predicted in all strains between - 515 and -471 nucleotides upstream of the initiator codon for 298 strains, and between -605 and - 570 for three strains (supp Figure 3).

Taken together, our data show that the highly conserved RlmQ methylase is playing important roles in human infections. Slower growth and increased biofilm formation are indicative of a chronic infection mode, and conversely, toxin production is a marker of acute infection. As RlmQ is associated with a better growth under physiological conditions, with an increase in cytotoxic activity, and a decrease in biofilm formation, we suggest that RlmQ might be important for the establishment of acute infection.

## CONCLUSION

We showed here that *S. aureus* RlmQ methylase, paralogous to *E. coli* RlmI, is the enzyme which methylates the position 7 of G2601 located in the helix H90 of 23S rRNA. Interestingly, the two paralogous proteins have evolved different activities in these taxonomically distant bacterial species, which show different modification patterns (in particular, G2574 in *E. coli* is not modified). Although the two enzymes act on two different modification sites, their target nucleotides are located in a large structured region of the 50S subunit, which was proposed to accommodate the transition of tRNAs during translation. Whether this divergent evolution between Gram-positive and Gram-negative bacteria may have consequences on ribosome functioning remained to be studied. Interestingly enough, we demonstrated that under conditions closer to infectious conditions, the *rlm*Q mutant strain displays clearly distinct phenotypes compared to the WT strain suggesting that RlmQ could play an important role during infection and/or colonization of the human host. Further experiments will be required to demonstrate if this is the modification of the 23S rRNA which contributes to these phenotypes or if RlmQ may have additional functions and RNA substrates in *S. aureus*. This study highlights that the functions of rRNA-specific modification enzymes expand to adaptive responses such as stress, environmental changes and during infections caused by pathogenic bacteria, a domain that needs to be exploited.

## MATERIAL AND METHODS

### Bacteria strains, plasmids, and growth conditions

The selected transposon mutants (NE553, NE1671, NE1442, NE189, NE807, NE1546 and NE1337) was obtained from the Nebraska mutant library produced by transposon *bursa aurealis* insertion (Fey et al. 2013). We also used 301 clinical strains isolated from patients suffering from 3 different types of infection as follows: 126 strains responsible for bacteraemia with or without endocarditis from the French national prospective multicenter cohort VIRSTA ((Le Moing et al. 2015), ENA accession numbers: PRJEB49354, PRJEB48298), 166 strains from the French severe community-acquired pneumonia cohort ((Gillet et al. 2021), ENA accession number: PRJEB54685) and 9 strains from necrotizing soft tissue infections (Baude et al. 2019).

Glycerol stocks from *S. aureus* USA300 (LAC), USA300 JE2, HG001, and the selected transposon mutants (NE553, NE1671, NE1442, NE189, NE807, NE1546 and NE1337) were streaked in BHI agar and growth overnight at 37°C. Starter cultures were prepared the following day by inoculating isolated colonies in 2 mL BHI medium and incubating them at 37°C with agitation at 180 rpm for 16 h. Overnight cultures were diluted to OD_600nm_ 0.05 in 20 mL BHI and grown to late exponential phase at 37°C. Cells were harvested by centrifugation at 2100 g for 10 min at 4°C and stored at - 20°C. Chloramphenicol (10 μg/mL) was added to medium when appropriate. Strains were also cultured in BHI broth at 37°C overnight, diluted at OD_600nm_ =0,05 in fresh medium (BHI or RPMI). 96-well plate was seeded with 200 ml and incubated at 37°C, 30°C or 42°C in TECAN plate reader for 24 h. OD at 600 nm was measured every 15 min after agitation.

Complementation of the *rlm*Q mutant (NE1546) was achieved by introducing the plasmid pCN38:wP:*rlm*Q:TT containing the gene *rlm*Q under a weak constitutive *bla*Z-derived promoter. Sequences of promoter and primers used for cloning are shown in **Supplementary Table S1**. The coding DNA sequence (CDS) and Shine-Dalgarno region of *rlm*Q was PCR amplified from *S. aureus* HG001 genomic DNA (HG001 and USA300 *rlm*Q CDS are identical) using primers *rlm*Q-F and *rlm*Q-R and cloned in the vector pEW (Menendez-Gil et al. 2020) between a modified *blaZ* promoter (wP) and the native *bla*Z transcriptional terminator (TT). Primers P*bla*Z-F and TT-rev were then used to amplify the region wP:*rlm*Q:TT from pEW:*rlm*Q and the resulting fragment was subcloned in vector pCN38 (Charpentier et al. 2004). Plasmid pCN38:wP:*rlm*Q:TT was electroporated into *S. aureus* RN4220 strain before being introduced in NE1546 strain as described elsewhere (Schneewind and Missiakas 2014) and transformants were selected with chloramphenicol. For complementation assays, the empty pCN38 plasmid was introduced in wild-type USA300 JE2 and NE1546.

### RNA extraction and purification of 23S ribosomal RNA

Total RNA from *S. aureus* was prepared as previously described (Antoine et al. 2019). 23S rRNA was purified from total RNA by size-exclusion chromatography using a Superose 6 10/300 column (GE Healthcare) eluted at 0.4 mL/min with 300 mM ammonium acetate. Fractions were analyzed in 1% agarose gel in TBE buffer and those containing pure 23S rRNA were pooled and concentrated by ethanol precipitation.

### AlkAniline treatment and primer extension

Detection of m^7^G was performed by *AlkAniline* treatment as described in (Marchand et al. 2018). Briefly, 5 μg of total RNA in 10 μL of water were mixed with 10 μL of 100 mM NaHCO_3_ (pH 9.2) and incubated at 96°C for 5 min. Alkaline hydrolysis was stopped by precipitating the RNA with 10 μL 3M sodium acetate (pH 5.2), 1 μL glycoblue and 1mL of ethanol overnight at -20°C. Treated RNA was pelleted by centrifugation (16000 g for 15 min at 4°C), washed with 70% ethanol, and dried under vacuum in a SpeedVac (Savant). RNA pellet was resuspended in 20 μL 1 M aniline (pH 4.2). and incubated at 60°C for 15 min. Aniline cleavage was stopped by precipitating RNA as above.

Analysis of 23S rRNA G2575 position in treated RNA was performed by primer extension using DNA primer H90 complementary to region 2645-2666 (**Supplementary Table S1**). 10 pmol of primer were ^32^P-5’-radiolabeled by incubation with γ-[^32^] ATP and 10 units of T4 polynucleotide kinase for 1 h at 37°C. Reverse transcription reactions were performed in a final volume of 10 μL. First, 100,000 cpm of labeled primer were incubated with 1 μg of treated RNA at 80°C for 90 s, then placed in ice for 90 s to allow hybridization. Two μL of reverse transcription buffer 5X were added and the samples were incubated at 20°C for 15 min before addition of 1.5 μL of dNTP mix (1 mM of dATP, dGTP, dCTP, dUTP) and 1 μL of AMV enzyme (PROMEGA). Reverse transcription was performed at 42°C for 45 min and synthesized cDNA was recovered by ethanol precipitation as described above. Dried pellets were resuspended in 6 μL urea-blue buffer (8 M urea, 0.025% xylene cyanol and bromophenol blue) and 30,000 cpm of each sample were separated by electrophoresis at 75 W in a denaturing 10% polyacrylamide-7 M urea gel until the bromophenol blue reached the bottom. The gel was then placed inside a cassette with an autoradiography (Kodak) and incubated overnight at -80°C.

### Mass spectrometry

LC/MSMS analysis was performed as previously described (Antoine et al. 2019). Modification at m^7^G2601 was identified by RNase T1 digestion of purified *S. aureus* 23S rRNA. The lack of methylation at C1989 was confirmed by *in-gel* RNase T1 digestion of a specific 23S rRNA fragment (nucleotides 1966 to 2068) isolated by RNase H cleavage using DNA primers H70-1 and H70-2 (**Supplementary Table S1**). RNase H reaction was performed in a final volume of 10 μL. First, 5 μg of pure 23S rRNA were incubated with 10 μg of primers at 80°C for 2 min and slowly cooled down to 50°C to allow hybridization. Then, 0.5 U of RNase H (Thermo Fisher Scientific) in the appropriate buffer were added, and digestion was for 30 min at 50°C. The RNase H fragment was isolated in a denaturing 10% polyacrylamide-urea gel and excised under UV light. RNase T1 digestion (20 μL) was performed with 20 U of enzyme in 100 mM ammonium acetate pH 6.8 at 50°C for 4 h. Samples were desalted using ZipTip C18 (Millipore) by several washes with 200 mM ammonium acetate, eluted in 50% acetonitrile and dry under vacuum. The pellet containing the RNase digest was resuspended in 3 μL of milli Q water and separated on a nanoAcquity UPLC system (Waters) equipped with a Acquity UPLC peptide BEH C18 column (130 Å, 1.7 μm, 75 μm × 200) equilibrated at 300 nL/min with a buffer containing 7.5 mM triethylammonium acetate, 7.0 mM triethylamine and 200 mM hexafluoroisopropanol. After loading, the oligoribonucleotides were eluted first with a gradient of methanol (15 – 35%) for 2 min, followed by another gradient of methanol (35 – 50%) for 20 min. Mass spectrometry (MS) and MS/MS analysis were performed using a SYNAPT G2-S instrument (Waters) equipped with a NanoLockSpray-ESI source in negative mode with a capillary voltage set to 2.6 kV and sample cone of 30 V. The source was heated to 130°C. Samples were analyzed over an *m/z* range from 500 to 1500 for the full scan, followed by a fast data direct acquisition scan (Fast DDA). Collision induced dissociation (CID) spectra were deconvoluted using MassLynx software (Waters) and manually sequenced by following the *y* and/or *c* series.

### Polysome profiling

*S. aureus* USA300 and NE1546 were cultivated in 20 mL BHI medium at 37°C until OD_600nm_ 2.0. Cells were resuspended in 500 μL lysis buffer (20 mM Tris pH 8, 100 mM NH_4_Cl, 50 mM MgCl_2_, 0.4% Triton X-100, 0.1% nonidet P-40), transferred into a tube with lysis matrix B (MP Biomedicals) and lyzed by bead beating in a FastPrep24 apparatus (MP Biomedicals) at 6 m/s for 40 sec. Cell debris were removed by centrifugation (16000 g for 10 min at 4°C) and the supernatant was recovered. Equal OD_260nm_ units of each sample were layered on top of 5-50% sucrose gradient prepared in buffer G (20 mM Tris pH 8, 100 mM NH_4_Cl, 15 mM MgCl_2_) and separated by ultracentrifugation in a Beckman SW-41Ti rotor at 39,000 rpm for 2 h 46 min at 4°C. Finally, samples were fractionated on a piston gradient fractionator (Biocomp) and the OD_260nm_ was recorded to generate sedimentation profiles.

### Eucaryotic cell and cultures conditions

Human monocytes (U937) expressing the C5a receptor were previously constructed (Spaan et al. 2013). This receptor is targeted by the Panton Valentine leucocidin (PVL) and the gamma toxin CB (HlgCB), two pore forming toxins of *S. aureus*. Thus *S. aureus* supernatant cytotoxicity on these cells is mainly driven by the activity of these two toxins.

### Biofilm formation

Strains were cultured in BHI at 37°C overnight, diluted at OD_600nm_ =0,01 in fresh medium. 96-well plate was seeded with 100 μL and incubated at 37°C in humidity chamber for 24 h. The mature biofilm was evaluated using crystal violet assay staining biomass. Then, 150 μL of crystal violet 1% was added for 15 min and washed using the steam technology (Tasse et al. 2018). The stain was solubilized for 15 min in 200 μL of 33% acetic acid and the OD_590nm_ was read in Tecan plate reader.

### Cytotoxicity assay

Bacterial strains were cultured in CCY broth at 37°C overnight, centrifuged at 10 000 g for 10 min and supernatant were collected. U937 cells were routinely cultured in DMEM growth medium supplemented with 10% foetal bovine serum at 37°C with 5% CO_2_. Cells were diluted at 1.10^6^ cell/ml and Iodure Propidium (IP) was added for a final concentration at 25 μg/mL. A 96-well plate was seeded with 90 μL of this solution and 10 μL of culture supernatant were added. IP incorporation into cells was measured using TECAN plate reader.

### Genomic analysis

All 301 clinical strains were previously sequenced, and all stages of quality control, sequence assembly and typing, as well as genes and non-coding RNA genes analysis were carried out (Baude et al. 2019; Gillet et al. 2021; Bastien et al. 2023). The GyrB, CoA and RlmQ sequences corresponding to SAUSA300_0005, SAUSA300_0224 and SAUSA300_0981 of the USA300_FPR3757 reference strain were compared with the 301 clinical strains. For each gene, a global alignment of the 301 sequences with the USA300_FPR3757 reference sequence was processed with clustal omega v1.2.4 (Sievers et al. 2011). A variation parameter λ = m/n where m represents the number of mutated amino acids and n the number of codons in the gene (Wagner 2007) was calculated. Promotech v1.0 software (Chevez-Guardado and Pena-Castillo 2021) was used in all 301 strains for promoter detection. DNA motif of the promoter was generated using MEME v4.11.2 (Bailey et al. 2009). Verification of the quality of the *rlm*Q mutant, in particular the absence of mutations outside the *rlm*Q locus, has been carried out. JE2 *rlm*Q^-^ transposon mutant genomic check was carried out using the JE2 strain as a comparator; validation of gene gain or loss was performed via a pangenome analysis and verified following the use of multiBamSummary from deeptools v3.5.0 to locate loci in the WT strain that were not covered by reads from the mutant strain. An additional analysis focusing on the presence of single nucleotide polymorphism (SNP) via variant calling was performed as in SNP analysis section (Bastien et al. 2023).

## ACKNOWLEDGEMENTS

This work was supported by the French National Research Agency ANR (SaRNAmod:ANR-21-CE12-0030-01 to SM; RHU IdBioriv: ANR-18-RHUS-0013 to FV), and by the Interdisciplinary Thematic Institute IMCBio+, as part of the ITI 2021-2028 program of the University of Strasbourg, CNRS and Inserm, IdEx Unistra (ANR-10-IDEX-0002), SFRI-STRAT’US (ANR 20-SFRI-0012), and EUR IMCBio (ANR-17-EURE-0023) under the framework of the French Investments of the France 2030 Program. B.P.K. acknowledges support from ANR, the epiRNA funding from the Region Grand Est, the FRISBI, Instruct-ERIC and iNEXT-Discovery infrastructures.

